# The Centroparietal Positivity Traces the Path of Evidence Accumulation During Dietary Decisions

**DOI:** 10.64898/2026.07.18.739320

**Authors:** Violet J. Chae, Lauren Fong, Tijl Grootswagers, Paul Garrett, Stefan Bode, Daniel Feuerriegel

## Abstract

Each day we make dozens of decisions about food. These choices are informed not only by information that we receive from our senses, but also by our memories, values, cultural norms, and goals. Evidence accumulation models have been proposed to explain how we transform these disparate sources of information into dietary choices and their corresponding motor actions. These models describe decisions arising from a noisy integration of decisional evidence until a criterion is reached. We investigated whether the centroparietal positivity (CPP), an event-related potential component proposed as a neural correlate of evidence accumulation during perceptual judgements, also traces accumulation trajectories during dietary decisions. Participants (*N* = 110) made speeded Yes/No judgements about food images relating to three attributes (healthiness, tastiness, and willingness to eat) while electroencephalography was recorded. They subsequently provided continuous ratings for each attribute. After applying signal deconvolution techniques, CPP waveforms exhibited a close correspondence to hypothesised evidence accumulation trajectories, showing steeper amplitude build-up rates for both faster decisions and continuous ratings more strongly in favour of one’s categorical choice. Our findings provide clear support for a neurally instantiated evidence accumulation process that integrates multiple sources of information to produce dietary decisions.

## Introduction

When faced with a restaurant menu, how do we decide what to eat? According to models of dietary decision-making (Rangel, 2013), we evaluate food items on relevant attributes to calculate an overall value for each option, and use these value representations to select the option with the higher overall value. Similarly, when confronted by a single food option – to eat or not eat the last slice of cheesecake – a decision may be reached by comparing the food’s representational value to an internal decision criterion.

Substantial progress has been made in understanding how our brains evaluate food items. Relevant food attributes (e.g., healthiness, tastiness, calorie content) are rapidly appraised (Chae et al., 2026; Meule et al., 2013; Moerel et al., 2024, 2026; Schubert et al., 2021; Toepel et al., 2009) and integrated into value signals in key regions, such as the ventromedial prefrontal cortex (Hare et al., 2009, 2011; Hutcherson et al., 2012). However, very little is known about how the brain transforms value representations into choices and actions. Understanding this process is essential for a complete account of how dietary decisions unfold in the brain.

In perceptual decision-making research, sequential sampling models characterise decision making as the accumulation of noisy evidence towards a choice and action outcome. The Diffusion Decision Model (DDM; Ratcliff, 1978; Ratcliff & Smith, 2004) is the most widely applied sequential sampling model and accounts for well-established behavioural phenomena, such as the speed-accuracy trade-off and the characteristic shapes of response time (RT) distributions in perceptual tasks (Ratcliff & McKoon, 2008; Ratcliff & Rouder, 1998). In the DDM, decisions arise from the noisy accumulation of evidence towards a response boundary. The drift rate describes the average rate and direction of evidence accumulation and reflects the combined strength of external sensory and internal mnemonic evidence (Ratcliff et al., 2016). During decision formation, accumulated evidence is continuously perturbed by stochastic noise, causing evidence to fluctuate towards or away from the response boundary. This means that on average, higher drift rates produce faster and more accurate responses, while within-decision noise produces trial-to-trial RT variability.

In humans, the centroparietal positivity (CPP), a positive-going event-related potential (ERP) component recorded using electroencephalography (EEG) over midline posterior scalp electrodes, has been identified as a neural signature of evidence accumulation in perceptual decision tasks (Kelly & O’Connell, 2013; O’Connell et al., 2012). The CPP exhibits two critical properties predicted by sequential sampling models. First, the amplitude build-up rate (waveform slope) prior to a motor response scales with RT and stimulus quality (steeper slopes for faster RTs and higher quality sensory evidence; Feuerriegel et al., 2021; Kelly & O’Connell, 2013; O’Connell et al., 2012; Twomey et al., 2015). Second, the amplitude just prior to the response remains approximately constant across RTs and stimulus quality conditions, consistent with accumulation to a fixed decision threshold (Kelly & O’Connell, 2013; O’Connell et al., 2012; Twomey et al., 2015). Establishing the CPP as a neural correlate of evidence accumulation provides a critical link between computational models and brain activity, revealing how the brain transforms sensory information into perceptual choices.

Sequential sampling models have been successful in accounting for speeded perceptual decisions based on sensory input, where the decision-relevant information is well defined (Bowman et al., 2012; Brunton et al., 2013; Ossmy et al., 2013; Ratcliff et al., 2009; van Vugt et al., 2012). These models have also been successfully applied to account for choice proportions and RT distributions in value-based decision tasks, such as choosing foods (Krajbich et al., 2010; Krajbich & Rangel, 2011; Milosavljevic et al., 2010) and consumer goods (Krajbich et al., 2012; Philiastides & Ratcliff, 2013), which additionally draw from endogenous sources of information (e.g., memories, knowledge, and personal preferences). This indicates that noisy evidence accumulation may serve as a general decision-making process that extends beyond perceptual choices (Kable & Glimcher, 2009).

Consistent with this idea, the CPP has been found to trace evidence accumulation dynamics during memory-based decisions (van Ede & Nobre, 2024), preference-based decisions (Fong et al., 2026), and crucially, two-option dietary decisions (Pisauro et al., 2017). Pisauro and colleagues presented participants with pairs of food items while recording EEG. They reported a centroparietal signal showing ramping-to-bound dynamics, with a build-up rate that was steeper for faster RTs and for trials with higher subjective value differences between the two foods. Taken together, these findings raise the possibility that evidence accumulation may be a domain-general decision-making mechanism indexed by a common neural marker, extending to dietary choice.

However, whether the CPP robustly reflects evidence accumulation during value-based and dietary decisions remains contested. Frömer et al. (2024) recorded EEG while participants made value-based choices between sets of consumer goods and food items. They found that, although a CPP-like ramping signal was observed, its properties did not align with the predictions of evidence accumulation models. The authors argued that the observed RT-dependent modulation of CPP slopes and waveform amplitudes can arise artifactually from the temporal overlap between stimulus-locked and response-locked EEG signals, rather than reflecting a genuine evidence accumulation signal. When they applied a deconvolution technique to separate these overlapping components, they could no longer detect waveform amplitude differences across fast and slow RT conditions that resemble CPP slope modulations. Frömer et al. also reanalysed EEG datasets from previously published studies, including the dietary choice study by Pisauro et al. (2017). By applying the same deconvolution procedure, they similarly attenuated or abolished waveform amplitude differences across fast and slow RT trials in those datasets.

While this study raised valid concerns regarding EEG signal overlap, the deconvolution-based analysis methods they used may not be suitable for identifying ramping evidence accumulation signals, and can artefactually prevent their detection (for a detailed critique see O’Connell et al., 2025). Resolving this debate requires studies that apply more appropriate methods to account for component overlap when characterising neural signals of evidence accumulation during dietary choice tasks.

An additional challenge when studying dietary decisions is that some valuation processes, such as attribute appraisal, may temporally overlap with choice-related processes, potentially hindering our ability to isolate the latter in the neural signals. When food images are presented, numerous attributes (e.g., healthiness, tastiness, edibility, familiarity) are rapidly represented by the brain with overlapping time-courses during the first second of viewing (Chae et al., 2026; Moerel et al., 2024; Schubert et al., 2021). In typical dietary and value-based decision paradigms, participants are presented with decision options (e.g., food images) and immediately make speeded choices (e.g., Frömer et al., 2024; Pisauro et al., 2017). This means that the appraisal of relevant attributes and the conversion of this information into action likely occurs in parallel, making it difficult to disentangle neural correlates of appraisal and choice.

One approach to better isolate evidence accumulation processes is to temporally separate attribute appraisal from choice, by presenting decision options first and then prompting a decision after a delay. In such a paradigm, the appraisal of each option can begin earlier and there is time to process decision-relevant attributes. The subsequent choice process draws primarily on endogenous memory representations rather than ongoing sensory processing. Previous work has used this approach to demonstrate that the CPP exhibits characteristic accumulation dynamics during working memory-based decisions (van Ede & Nobre, 2024). However, they did not use signal deconvolution to account for choice prompt stimulus-locked EEG signals, which largely overlapped in time with their response-locked waveforms.

We investigated whether the CPP exhibits signatures of evidence accumulation during dietary decisions, extending this framework beyond perceptual choices. Participants viewed food images and gave binary (Yes/No) responses according to whether the food was healthy, tasty, or whether they were willing to eat it, while EEG was recorded. These decisions (single-item choice and healthiness/tastiness judgements) differ from the two-choice food preference decisions studied previously (Pisauro et al., 2017). Nevertheless, such choices are frequently made in everyday life and may recruit domain-general evidence accumulation mechanisms. Our paradigm separated attribute appraisal processes from the subsequent conversion of this information into choice. We first presented the food image in isolation and then prompted participants to make one of the three possible categorisation decisions after a delay (Figure 1A). This design allowed us to better isolate the neural mechanisms underlying the processes leading up to the decision and corresponding motor action. After applying an appropriate deconvolution method to account for EEG signals time-locked to stimulus onset, we assessed whether the CPP consistently displayed the hallmark characteristics of evidence accumulation across these three types of dietary judgements.

**Figure 1.**
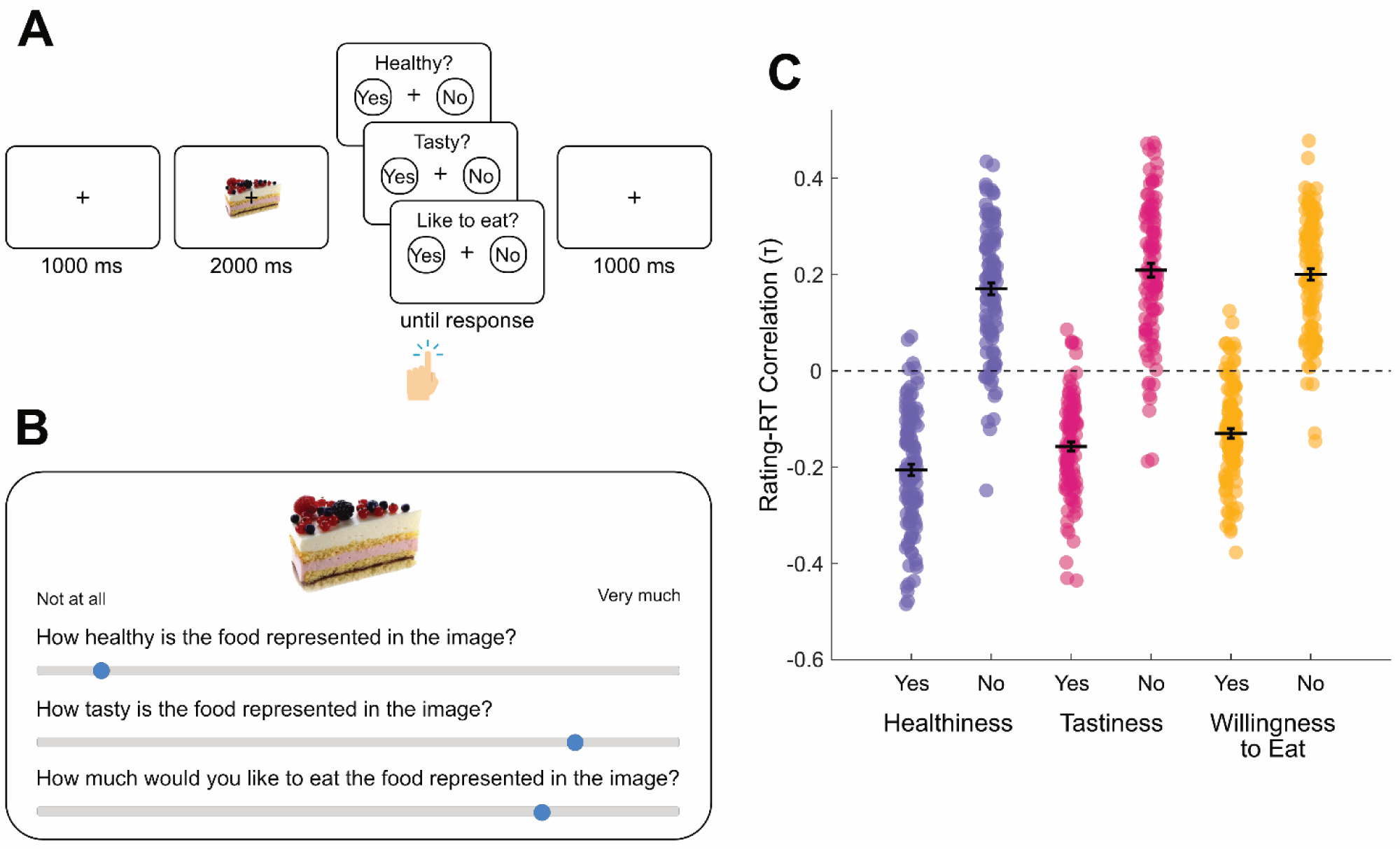
Trial diagrams and correlations between continuous ratings and categorisation task RTs. A) EEG food categorisation task. In each trial, participants viewed a food image (2 s), followed by a response screen displaying one of three decision prompts (healthiness, tastiness, or willingness to eat) and one of two keyboard press response mappings (‘z’ key for Yes and ‘m’ key for No, or the reverse). Participants were instructed to respond as quickly as possible. B) Continuous rating task. Participants provided continuous ratings for each food image for the three attributes (healthiness, tastiness, willingness to eat) using sliding scales ranging from 0 (“Not at all”) to 100 (“Very much”). C) Participants’ rating-RT correlations. Kendall’s tau-b correlation coefficients between continuous ratings and RTs for Yes and No trials separately within the three decision types, shown for individual participants (dots) and group means (horizontal black lines). Error bars denote standard errors.

## Methods

The experiment consisted of two sessions. In the first session, participants completed a food categorisation task while EEG was recorded and subsequently provided food attribute ratings for each food image. In the second session, participants completed two behavioural food choice tasks and answered questionnaires regarding their dietary style and food motivations. Data collected from the second session were not analysed in this study, and the behavioural tasks and the questionnaires are described in Chae et al. (2025). This study was approved by the Human Research Ethics Committee of the University of Melbourne (HREC ID 24850).

### Participants

Participants (*N* = 117) were recruited through the University of Melbourne student portal and advertising via posters. Participants indicated that they were at least 18 years old, were fluent in written and spoken English, did not have a history of any eating disorders, were not on a calorie restriction diet, and did not have any dietary restrictions (e.g., for health or religious reasons). Participants were asked to fast for three hours prior to the start of both in-person sessions. At the end of the second session, participants were reimbursed AUD $50. The sample had a mean age of 26.8 years (SD = 8.50, range 18-57 years). 84 participants identified as female, 32 as male, and one as other.

### Procedure

Participants completed a food categorisation task presented using Psychtoolbox (Kleiner et al., 2007) interfacing Matlab R2022a. After the task, participants rated each food image on healthiness, tastiness and how much they would like to eat the food (willingness to eat) using Qualtrics.

#### Food Categorisation Task

Participants viewed each food item three times: once each across the healthiness, tastiness, and willingness to eat trials (Figure 1A). Food images were presented centrally, subtending 6.6 × 3.5 degrees of visual angle on a 24.5-inch ASUS ROG Swift PG258Q monitor (1920 x 1080 pixels, 60 Hz refresh rate). Participants’ heads were stabilised using a chin and forehead rest approximately 50 cm from the monitor.

Each trial began with a fixation cross presented in isolation for 1 s, followed by the food image with a fixation cross for 2 s. Next, the participants were prompted with “Healthy?” in a healthiness trial, “Tasty?” in a tastiness trial, and “Like to eat?” in a willingness to eat trial, along with a response mapping that indicated which of the left (“z”) and the right (“m”) keyboard keys corresponded to Yes and No responses. There was an equal chance of the left keyboard press indicating a Yes response (and the right keyboard press indicating a No response) and the reverse for each trial. For each participant, the order of healthiness, tastiness, and willingness to eat trials were randomised. The randomisation of the trial type order and response mapping ensured that participants could not pre-emptively prepare a keypress motor action during the food image presentation period.

Participants were asked to respond as quickly and accurately as possible and to pay attention to the switching of the response mapping across trials. Participants completed five practice trials and indicated that they had understood the instructions before starting the main experiment. To reduce eye movement artifacts the participants were instructed to keep their eyes fixated on the fixation cross, which remained on screen throughout the duration of the experimental blocks. Response time was recorded as the time taken from the presentation of the prompt to the participant’s response via keyboard press. We randomised image presentation order for each participant (while ensuring that the same food image was not presented in two consecutive trials).

#### Continuous Ratings

Participants rated each food image on healthiness (“How healthy is this food?”), tastiness (“How tasty is this food?”) and how willing they were to eat it (“How much would you like to eat this food?”). Participants were asked to use a computer mouse to indicate their response on a sliding scale from “Not at all” to “Very much” (Figure 1B). The order of image presentation was randomised.

### Behavioural Data Analyses

To verify whether choices and RTs in the categorisation task reflected stable judgments of each food attribute, we assessed correlations between RTs and the continuous ratings provided for each food, for Yes and No choices separately. Kendall’s tau-b correlations were chosen to account for rank ties in the continuous ratings. Correlation coefficients were obtained for each participant and attribute and are plotted in Figure 1C.

### EEG Recording s Data Processing

EEG was recorded using a Biosemi Active II system (Biosemi, The Netherlands) with 64 channels at a sampling rate of 512 Hz using common mode sense and driven right leg electrodes (http://www.biosemi.com/faq/cms&drl.htm). We attached 64 electrodes to the cap according to the international 10-20 system. We also attached eight additional electrodes: two electrodes placed 1 cm from the outer canthi of each eye, four electrodes placed above and below the centre of each eye, and two electrodes placed above the left and right mastoids. Electrode offsets were kept within the range of ±20 µV during recording.

EEG data was processed using EEGLab v2022.1 (Delorme & Makeig, 2004) interfacing Matlab (R2022a). Excessively noisy channels were identified through visual inspection (mean number of bad channels = 0.26, range 0–5) and excluded from average reference calculations and independent components analysis (ICA). Excessively noisy sections or those with large amplitude artefacts were identified through visual inspection and removed. The data were referenced to the average of all channels (excluding excessively noisy channels). One channel (AFz) was removed to compensate for the rank deficiency due to average referencing. EEG data were low-pass filtered at 30 Hz (EEGLab Basic Finite Impulse Response Filter New, default settings). A copy of the dataset was created for the purpose of ICA. A 0.1 Hz high-pass filter (EEGLab Basic Finite Impulse Response Filter New, default settings) was applied to the copied dataset to improve stationarity for the ICA. We used RunICA extended algorithm (Jung et al., 2000) to perform the ICA. The resulting independent component information was copied to the original dataset (e.g., as done by den Ouden et al., 2023). Independent components associated with eye blinks and saccades were identified and removed in line with guidelines in (Chaumon et al., 2015). Next, we interpolated the excessively noisy channels and AFz using spherical spline interpolation.

EEG data were initially segmented from -1300 to 4200 ms relative to food image onset and baseline corrected using the data from the 200 ms time window prior to the food image onset. Epochs containing amplitudes exceeding ±200 µV at any scalp channel were excluded from the analyses.

#### EEG Signal Deconvolution

In tasks where stimuli and responses are separated by variable intervals, ERP components time-locked to stimulus onset can overlap temporally with response-locked neural activity, potentially confounding estimates of pre-response signals such as the CPP. To mitigate this, we applied Residue Iteration Decomposition (RIDE; Ouyang et al., 2015) to separate stimulus-locked contributions from response-locked and variable latency EEG signals. RIDE decomposes single-trial EEG data into distinct subcomponents based on their temporal alignment to different events within each epoch, making it suitable for recovering latency-variable and response-locked signals (Ouyang et al., 2015). This approach addresses concerns raised by Frömer et al. (2024) regarding artefactual RT-dependent CPP slope differences arising from overlapping stimulus- and response-locked activity, and has been validated for use with ramping evidence accumulation signals in prior work (Fong et al., 2026; Steinemann et al., 2018).

We decomposed the stimulus-locked single-trial EEG data for each participant into three subcomponents using the following time windows: a stimulus-locked (S) subcomponent estimated from 0 to 1000 ms post-stimulus onset, a response-locked (R) subcomponent estimated from −600 to −100 ms relative to response onset, and a time-varying (C) subcomponent estimated from 500 to 1000 ms post-stimulus, which captures neural activity with a latency that varies across trials and is not strictly tied to either the stimulus or response. We took a conservative approach to estimating and removing stimulus-locked EEG activity, using an extended estimation window for the S subcomponent beyond the default recommendations of Ouyang et al. (2015), following Sun et al. (2024) and Fong et al. (2026). This was done to minimise the risk that response-locked signals reflecting genuine decision processes, such as evidence accumulation, might be confounded by residual stimulus-evoked activity. The estimated S subcomponent was subsequently subtracted from each stimulus-locked single trial epoch prior to extracting stimulus- and response-locked epochs, such that the resulting waveforms more selectively capture time-varying and response-locked neural dynamics underlying decision formation. R and & subcomponent estimates were included to improve decomposition accuracy in our estimation of the S subcomponent but were not otherwise used in subsequent analyses.

Following S subcomponent subtraction, we created response-locked epochs spanning −1300 to 200 ms relative to response onset. Epochs were baseline-corrected using the 100 ms window preceding food image onset, preserving the pre-stimulus baseline for response-locked ERP analyses.

#### Current Source Density Transformation

After deriving response-locked epochs, we applied a current source density (CSD) transformation to the response-locked data using the CSD Toolbox (Kayser & Tenke, 2006; m-constant = 4, λ = 0.00001 following Feuerriegel et al., 2021; Fong et al., 2026). CSD transformation estimates the second spatial derivative of the scalp-recorded potential, enhancing the spatial resolution of the EEG signal by emphasising local cortical sources and attenuating the contribution of spatially diffuse signals. This step was particularly important for isolating the CPP measured at posterior electrodes from co-occurring signals generated at more anterior frontal and central sites (e.g., motor preparatory activity) which can spread to parietal electrodes via volume conduction (Feuerriegel et al., 2022; Fong et al., 2026; Kelly & O’Connell, 2013).

#### Event-Related Potential Analyses

We examined the relationship between the pre-response CPP component and both RTs and continuous attribute ratings using two complementary approaches: analyses focused on established pre-response analysis windows, and mass-univariate analyses spanning the full response-locked epoch.

#### Deriving CPP Measures

Pre-response CPP slope and amplitude measures were extracted from response-locked epochs at electrode Pz. Slopes were estimated by fitting a linear regression to each single-trial waveform within a window spanning −300 to −70 ms relative to response onset. Pre-response amplitude was defined as the mean signal within −130 to −70 ms relative to response onset. This time window was chosen to capture the amplitude of the CPP around the onset time of motor execution prior to completion of the keypress (in line with previous work, e.g., Feuerriegel et al., 2022; Fong et al., 2026; Steinemann et al., 2018).

### Window-Based Analyses Involving RT

We used linear mixed effects models to test whether RT predicted CPP slope and amplitude. Models were fit separately for each decision type (healthiness, tastiness, and willingness to eat). Prior to modelling, RTs were log-transformed to reduce positive skew and then z-scored within each participant to remove between-participant differences in RT scale, allowing inference to focus on within-participant variance (as done by Fong et al., 2026). We specified a maximal random-effects structure that included participant random intercepts and random slopes for the predictor of interest:

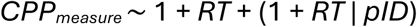

Where the maximal model produced a singular fit or convergence failure (e.g., due to insufficient trial counts, near-zero variance components, or collinearity), the random slope was removed and only a random intercept was retained (Bates et al., 2015). We compared models with and without the fixed effect of RT but with identical random effects structures using likelihood ratio tests under maximum likelihood estimation. We report regression coefficients (β) from the best-fitting model that converged without singularity. Full likelihood ratio test statistics and regression coefficients are reported in Supplementary Tables 2 and 3, respectively.

### Window-Based Analyses Involving Continuous Attribute Ratings

We assessed whether participants’ attribute ratings correspond to differences in subjective attribute values and CPP slope, analogous to the rate of decisional evidence accumulation (drift rate) in sequential sampling models. We used rank correlation analyses (Kendall’s tau-b) to test for ordinal relationships between the pre-response CPP measures and continuous attribute ratings. Correlation analyses were run for Yes and No trials separately for each decision type (healthiness, tastiness, willingness to eat). Fisher-transformed correlation coefficients were tested against zero at the group level using one-sample t-tests (two-tailed). Note that there were more Yes trials than for No trials for most participants, particularly for tastiness and willingness to eat decisions. The descriptive statistics for the trial counts in each condition are reported in the Results. If there were less than 20 trials available for the correlation analyses, the participant was excluded from further analyses for that decision type. All participants (*N* = 110) were retained for Yes trials for all decision types. We retained 109 participants for healthiness No trials, 105 participants for tastiness No trials, and 106 participants for willingness to eat No trials.

### Mass-Univariate Analyses for RT and Continuous Ratings

To examine the relationships between RT/continuous ratings and CPP measures beyond the fixed analysis windows, we performed complementary mass-univariate analyses to test for associations across the full response-locked epoch. At each timepoint, CPP slope was estimated as the linear slope of a 300 ms window centred on that timepoint; timepoints within 150 ms of the epoch boundaries were excluded as a full 300 ms window could not be constructed. Amplitude at each timepoint was taken as the instantaneous signal value.

For RT, we fit linear regression models at each timepoint to predict the CPP slope and amplitude values using RT as the predictor, yielding time series of β coefficients. This was carried out separately for each decision type (healthiness, tastiness, willingness to eat). One-sample t-tests were performed to compare the distributions of the resulting β values against 0. To correct for multiple comparisons, we used cluster-based permutation tests (10,000 permutations, cluster-forming threshold: *p* < .01), implemented in the Decision Decoding Toolbox (Bode et al., 2019; Maris & Oostenveld, 2007). Significant clusters were identified as those whose summed t-value exceeded the 97.5^th^ percentile of the permutation-derived null distribution (α = .05, two-tailed). For continuous ratings, we calculated the Kendall’s rank correlation coefficient (tau-b) at each timepoint to between the CPP slope and amplitude values and continuous ratings, yielding time series of τ coefficients. This was carried out for Yes and No trials separately for each decision type (healthiness, tastiness, willingness to eat). Correlation coefficients were Fisher transformed and one-sample, single-tailed t-tests were performed to compare whether the distributions of the transformed τ values were above zero (Yes trials) or below zero (No trials). To correct for multiple comparisons, we used cluster-based permutation tests as outlined above.

Bayes factors (BF10) were also computed for each timepoint using the BayesFactor toolbox v3.0 (Krekelberg, 2024) using default Cauchy priors (r = 0.707; Rouder et al., 2017). Bayes factors are reported here solely as a descriptive index showing where evidence supports the presence versus absence of an effect at each time point.

## Results

Participants (*N* = 110) completed a food categorisation task in which they viewed food images and made speeded binary (Yes/No) choices relating to one of three food attributes (healthiness, tastiness, and willingness to eat; Figure 1A). Participants indicated their choice via keypress with their left or right index finger and EEG was recorded throughout the task. In a separate phase conducted without EEG recording, participants rated each food image on the same three attributes on a continuous sliding scale, with 0 indicating “Not at all” and 100 indicating “Very much” (Figure 1B).

We first assessed whether categorisation task performance was associated with the subjective continuous attribute ratings. Participants responded faster for foods with higher ratings (closer to “Very much”) for Yes trials (healthiness: mean τ = -.21, *SD* = .13, *t*(109) = -17.32, *p* < .001, tastiness: mean τ = -.16, *SD* = .10, *t*(109) = -16.23, *p* < .001, willingness to eat: mean τ = -.13, *SD* = .10, *t*(109) = -13.55, *p* < .001) and for foods with lower ratings (closer to “Not at all”) for No trials (healthiness: mean τ = .17, *SD* = .13, *t*(108) = 13.54, *p* < .001, tastiness: mean τ = .01, *SD* = .09, *t*(104) = 15.03, *p* < .001, willingness to eat: mean τ = .02, *SD* = .09, *t*(105) = 16.83, *p* < .001) for all three decision types (Figure 1C). These results demonstrate that the ratings reflect stable food evaluations and that the speed of participants’ choices reflected the strength of their internal representations for each attribute, validating our approach of linking retrospectively collected ratings to trial-level neural activity.

### CPP Morphology is Consistent with Evidence Accumulation Dynamics During Dietary Decision-Making

The CPP is defined as an ERP component with a clear parietal locus and a ramping-to-threshold signal morphology. We applied current source density (CSD) transformation, a spatial filtering technique that enhances the spatial resolution of EEG signals (Kayser & Tenke, 2006). This was done to better isolate signals at parietal electrodes and disambiguate them from concurrent EEG signals at more anterior and posterior sites, which can bias CPP waveform measurements (as done by Feuerriegel et al., 2021, 2022; Fong et al., 2026; Kelly et al., 2021; Kelly & O’Connell, 2013; Steinemann et al., 2018). We also applied a signal deconvolution technique (RIDE; Ouyang et al., 2015) to account for effects of stimulus-locked EEG signals (i.e., locked to decision prompt onset) that may temporally overlap with ramping evidence accumulation signals (Fong et al., 2026; Steinemann et al., 2018). Early sensory evoked EEG components may overlap more substantially with signals of interest for trials with faster as compared to slower RTs (Frömer et al., 2024), leading to spurious findings of a ramping-to-bound morphology (but see O’Connell et al., 2025). We used RIDE to isolate the stimulus-locked component of the EEG signal for each decision type separately. We then subtracted this stimulus-locked component from the single trials of EEG data for that decision type to better isolate time-varying, decision-related EEG activity such as ramping evidence accumulation signals.

ERPs at parietal electrode Pz, shown in Figure 2, were highly consistent with a ramping-to-threshold morphology. As visible in Figure 2C, ERP amplitudes immediately prior to the motor response showed a clear parietal positivity for all three decision types, consistent with the observed topography of the CPP in CSD-transformed data (Kelly & O’Connell, 2013; Feuerriegel et al., 2021; Fong et al., 2026).

**Figure 2.**
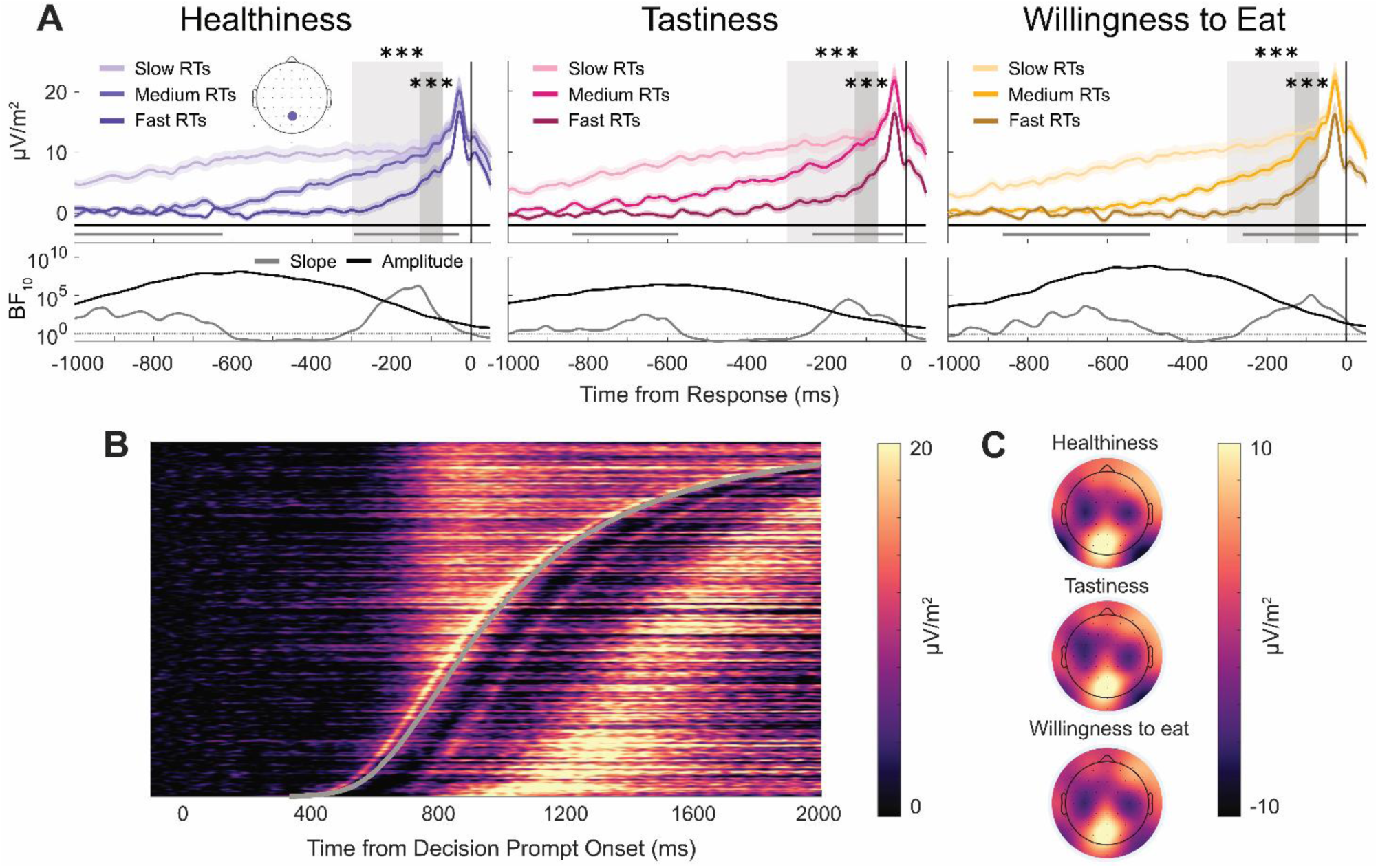
Associations between RT and CPP slope and amplitude for food-related decisions. A) Group-averaged response-locked ERP waveforms for healthiness, tastiness, and willingness to eat decisions, plotted by RT tertile (fast/medium/slow). Shaded areas denote standard errors. Pre-response analysis windows are denoted with light grey shading for slope analyses (−300 ms to −70 ms relative to response onset) and dark grey shading for amplitude analyses (−130 ms to −70 ms). Asterisks indicate statistically significant associations between RT and CPP slope and amplitude within the respective pre-response windows (*** indicates *p* < .001). Horizontal grey and black lines in the upper plots indicate the time windows with statistically significant associations between RT and CPP slope and amplitude respectively, derived from mass-univariate analyses. In the lower plots, grey and black lines indicate Bayes Factors (BFs) at each time point for associations between RT and CPP slope and amplitude respectively. B) Single-trial ERP amplitudes locked to decision prompt onset for all decision types, pooled across all participants and sorted by RT (fast RT below). Data were smoothed over 300 trial bins using a sliding Gaussian-weighted moving average filter. RTs are indicated by the grey line. C) Scalp topographies for healthiness, tastiness, and willingness to eat decisions, averaged across a pre-response analysis window (-130 ms to -70 ms relative to response onset).

Consistent with a ramping evidence accumulation signal, we found that CPP slopes across several hundred milliseconds leading up to the response were steeper for trials with faster decisions. Figure 2A shows the group-averaged response-locked waveforms for the three decision types, split into RT tertiles (fast, medium, and slow) for each participant and averaged across participants. Faster RTs were associated with steeper slopes across all decision types (healthiness: β = −0.02, *SE* = 0.003, *t*(104.55) = −5.68, *p* < .001, tastiness: β = −0.01, *SE* = 0.003, *t*(102.57) = −3.88, *p* < .001, willingness to eat: β = −0.01, *SE* = 0.003, *t*(12538.33) = −4.97, *p* < .001).

Outside of the pre-response analysis window, we found additional evidence for effects of RT on ERP slopes that were consistent with a ramping-to-bound morphology. We performed mass-univariate analyses to examine associations between CPP slope and RT across the full response-locked epoch. For each decision type, we identified two windows in which RTs were associated with CPP slopes, as denoted by horizontal grey lines in Figure 2A. In the earlier window, slower RTs were associated with steeper CPP slopes, consistent with a ramping evidence accumulation signal starting further back in time from the response in trials with slower decisions (from -1000 ms to -629 ms for healthiness, from -838 ms to -574 ms for tastiness, and from -863 ms to -492 ms for willingness to eat decisions). By contrast, faster RTs were associated with steeper CPP slopes in a later window that overlapped with the pre-response analysis window, extending to around the time of the response (from -295 ms to -29 ms for healthiness, from -235 ms to -8 ms for tastiness, and from -260 ms to +29 ms for willingness to eat decisions).

To further visualise the CPP waveform dynamics, we generated a heatmap of single-trial ERPs locked to decision prompt onset at electrode Pz (Figure 2B). Trials were sorted by RT (faster trials at the bottom) and were pooled across participants. As all decision types showed similar ERP dynamics and comparable pre-response amplitudes, we also pooled data across decision types. Consistent with gradual evidence accumulation towards a decision threshold (Fong et al., 2026; O’Connell et al., 2012), the heatmap reveals that CPP amplitudes build gradually with similar onset latencies across trials, and then converge to comparable amplitudes near the time of the response (curved grey line).

If the CPP traces evidence accumulation to a fixed criterion, pre-response amplitudes should not substantially vary across RTs, as all decisions terminate at the same threshold regardless of accumulation speed. However, we observed that faster RTs were associated with lower amplitudes consistently across all three decision types. Faster RTs were associated with lower amplitudes for all decision types (healthiness: β = 2.14, *SE* = 0.63, *t*(111.09) = 3.41, *p* < .001, tastiness: β = 2.85, *SE* = 0.80, *t*(110.83) = 3.57, *p* < .001, willingness to eat: β = 3.26, *SE* = 0.79, *t*(109.52) = 4.12, *p* < .001). As visible in Figure 2A, pre-response CPP amplitudes were attenuated in trials with fast RTs, visible as lower amplitudes for the fast RT tertile bin. By comparison, there were no clearly visible differences between the medium RT and the slow RT tertile bins. Plots of amplitudes by RT using finer-grained quantile bins (displayed in Supplementary Figure 1) revealed that amplitudes were lowest for trials with the fastest RTs and gradually increased across quantiles with longer RTs up to one-second, after which amplitudes appeared to be relatively stable.

Follow-up mass-univariate analyses similarly revealed that faster RTs were associated with lower amplitudes for the full second leading up to the time of response, including the pre-response analysis window, for all three decision types (black horizontal lines in the upper plots in Figure 2A; cluster *p* < .001 for healthiness, tastiness, and willingness to eat decisions). While earlier amplitude differences are expected due to the differences in CPP build-up rates, amplitude modulations immediately prior to the response are not in line with evidence accumulation to a fixed threshold.

We suspected that this amplitude decrement for the fastest trials may be due to some proportion of choices made prematurely, where computer keys were pressed without a corresponding evidence accumulation process that would be comparable to decisions in slower RT trials. These fast contaminant responses are well-documented in perceptual decision-making research (Ratcliff & Kang, 2021; Ratcliff & Tuerlinckx, 2002) and can arise due to a build-up of preparatory motor activity that precedes the onset of evidence accumulation, also resulting in reduced pre-response CPP amplitudes for very fast RTs (Kelly et al., 2021). We reasoned that if such responses were prevalent in fast RT trials, then there would be a substantially weaker correspondence between the continuous attribute ratings and categorical choices as compared to slower RT trials. This is analogous to observations of low accuracy in perceptual decision tasks for trials with very fast RTs (e.g., Kelly et al., 2021; Ratcliff & Kang, 2021).

To examine this possibility, we calculated differences in continuous attribute rating scores across Yes and No trials for each attribute across faster and slower RT quantiles (shown in Supplementary Figure 2). Large average rating differences between Yes and No trials indicate a high degree of choice consistency. Lower rating differences indicate a higher proportion of random responding, similar to low-accuracy, very fast RT trials in perceptual tasks. On average, we did not observe a substantial drop in continuous rating differences for the fastest RT trials, indicating that such contaminant responses were not prevalent in our data, and that even very fast choices were generally congruent with participants’ continuous attribute ratings.

By also performing additional simulation analyses (detailed in Supplementary Figure 3), we found that the amplitude differences by RT may be attributable to the deconvolution procedure used to isolate the CPP from overlapping stimulus-locked EEG signals. The CPP is a ramping signal that begins a short period after stimulus onset (O’Connell et al., 2025). Consequently, deconvolution procedures such as RIDE may erroneously attribute a portion of the ramping signal to the stimulus-locked component of the data, which is subtracted from single trials of EEG data prior to analyses (for examples of comparable artefacts see Frömer et al., 2024; O’Connell et al., 2025). By simulating stimulus-locked and ramping EEG signals and applying RIDE, we observed this misestimation of stimulus-locked signals.

As the signals captured by the stimulus-locked component fall closer to the time of the response for faster RT trials, the reduction in waveform amplitudes around the time of the response is expected to be greater in those trials. As this attenuation disproportionately affects faster responses, it may artificially generate the observed modulation of pre-response amplitude by RT. This pattern was also visible in our simulations, where amplitudes around the response were disproportionately attenuated for fast RT trials (Supplementary Figure 5). Consistent with this, the RT-amplitude association in our EEG data appears to be mostly driven by a drop in amplitude for the fastest trials (dark coloured lines; Figure 2A) rather than a general, linear difference in amplitude across RTs (similar peaks for medium and lighter coloured lines; Figure 2A). In addition, the time-course of pre-response CPP amplitude decrements observed in our data closely mirrored the amplitude profile of the RIDE-estimated stimulus-locked component (Supplementary Figure 4).

In summary, while the RIDE deconvolution procedure appeared to recover differences in ramping accumulator signals well, and is required to account for biasing effects of stimulus-locked ERPs, the biases observed in the simulations provide a plausible reason for the amplitude differences in our data.

### CPP Build-Up Rates Covary with Continuous Attribute Ratings

We also tested whether the CPP displayed build-up rates that were modulated by subjective attribute values, estimated using the continuous attribute ratings. This approach parallels how stimulus quality is operationalised in perceptual tasks. Just as higher motion coherence in random dot motion tasks provides higher stimulus quality for a direction judgement and produces higher evidence accumulation rates (drift rates; Kelly & O’Connell, 2013), more extreme attribute ratings indicate stronger representations for categorical judgements about a given food and correspondingly higher drift rates. For foods categorised as Yes, higher ratings indicate relatively stronger evidence favouring the Yes response, whereas for foods categorised as No, lower ratings indicate stronger evidence favouring the No response. Ratings further from the participant’s subjective category boundary should correspond to higher drift rates and steeper CPP build-up.

Examination of the RT distributions across continuous rating tertiles (Figure 3) revealed patterns consistent with differences in drift rates. The fastest trials (10^th^ RT percentiles) did not differ substantially between high, medium, and low continuous ratings. On the other hand, the slowest trials (90^th^ RT percentiles) exhibited the greatest rating-dependent variation. The influence of ratings on RT increased across faster to slower quantiles, consistent with trial-level changes in drift rate (Ratcliff & McKoon, 2008). Stronger representations (indexed by higher ratings for Yes trials and lower ratings for No trials) produce faster evidence accumulation, an effect that compounds over time and becomes most pronounced in the slowest responses within each rating tertile.

**Figure 3.**
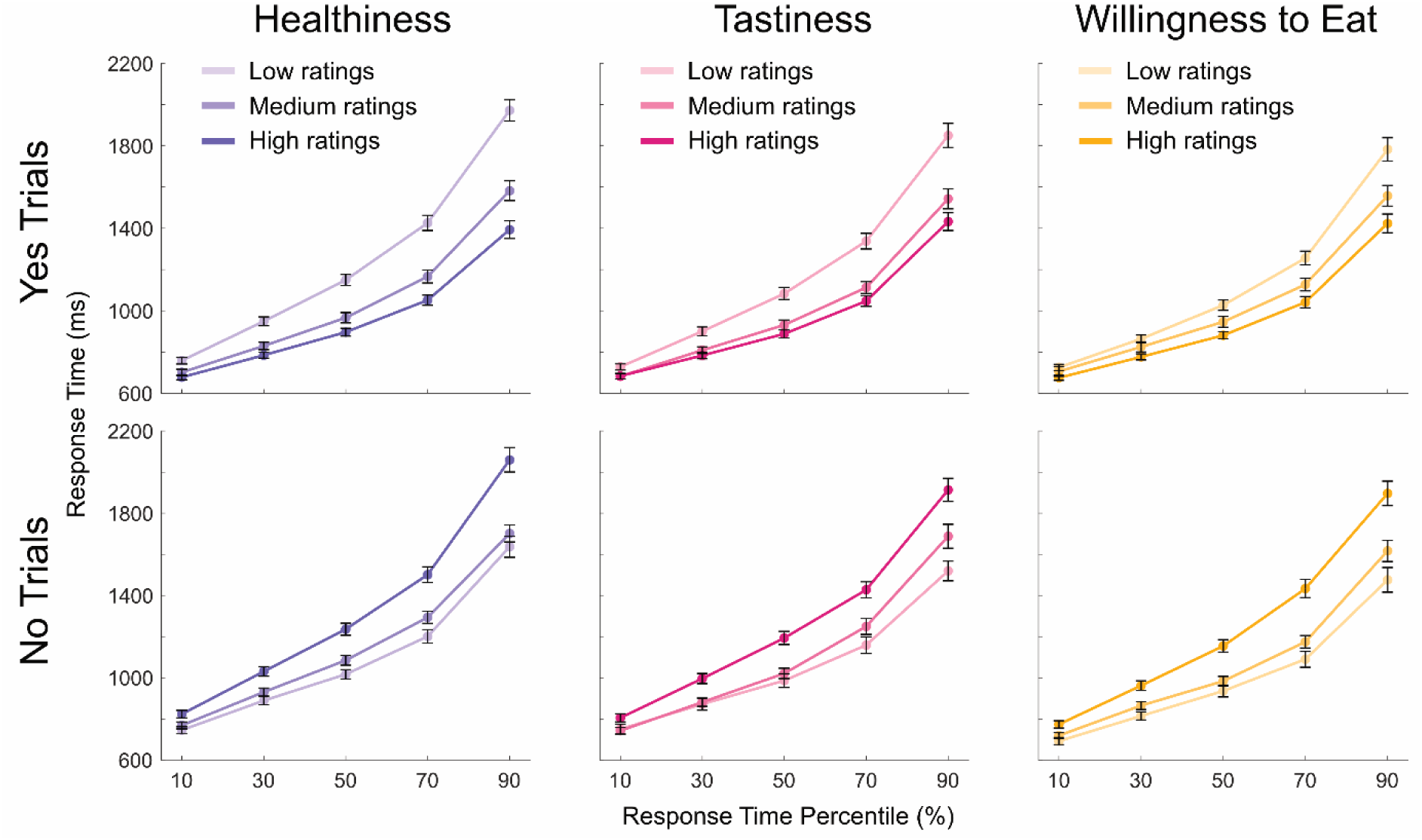
RT distributions stratified by continuous ratings. RT distributions at 10^th^, 30^th^, 50^th^, 70^th^, and 90^th^ percentiles for continuous rating tertiles (low, medium, and high), for Yes and No trials. Continuous ratings were divided into tertiles for each participant and decision type, then RT percentiles were calculated from the corresponding trial subsets before averaging across participants. Coloured circles represent group mean RTs for each percentile. Black error bars denote standard errors. The increasing separation between rating tertiles from fast to slow RT percentiles reveals that subjective attribute value (indexed by continuous ratings) has minimal influence on rapid decisions but substantial influence on slower decisions, consistent with drift rate modulation in evidence accumulation models.

Assuming that continuous ratings are correlated with single-trial drift rates, we expected to observe steeper pre-response CPP slopes in trials with higher ratings for Yes trials, and steeper slopes in trials with lower ratings for No trials, for all decision types. Importantly, we expected these associations to be more subtle than those between RT and CPP slope. The CPP is described as tracing the path of evidence accumulation in each trial, which includes both the influence of systematic drift and within-trial decision noise as specified in the DDM. The noise component substantially contributes to RT variability, so that even when drift rates differ on average, there can be substantial overlap in RT distributions and single-trial evidence accumulation profiles across lower and higher drift rate conditions, reflected in the RT percentile plots in Figure 3. In our data, there is high RT variability within each rating tertile but relatively small average RT differences across tertiles (in contrast to Kelly & O’Connell, 2013). In addition, the continuous ratings are unlikely to perfectly correspond to drift rates during the categorisation task due to changes-of-mind or decision noise influencing choices/ratings in each task, and we cannot reasonably assume a linear linking function between rating values and drift rates. Our large sample allowed us to test for these subtle associations using ordinal rank correlation analyses.

Figure 4 shows group-averaged response-locked waveforms for Yes and No trials for the three decision types, split by continuous rating tertile (high, medium, and low) for each participant and averaged across participants. Participants’ ratings were positively correlated with pre-response CPP slopes for Yes trials for all decision types (healthiness: mean τ = .04, *SD* = .09, *t*(109) = 4.25, *p* < .001, tastiness: mean τ = .01, *SD* = .09, *t*(109) = 1.71, *p* = .045, willingness to eat: mean τ = .02, *SD* = .09, *t*(109) = 2.94, *p* = .002). As expected, these associations were more subtle than those between CPP slope and RT, as visible in Figure 4. Ratings were negatively correlated with CPP slopes for No trials for healthiness decisions (mean τ = -.02, *SD* = .13, *t*(108) = -1.71, *p* = .045). However, we did not find clear evidence that ratings were associated with CPP slopes for No trials for tastiness and willingness to eat decisions.

**Figure 4.**
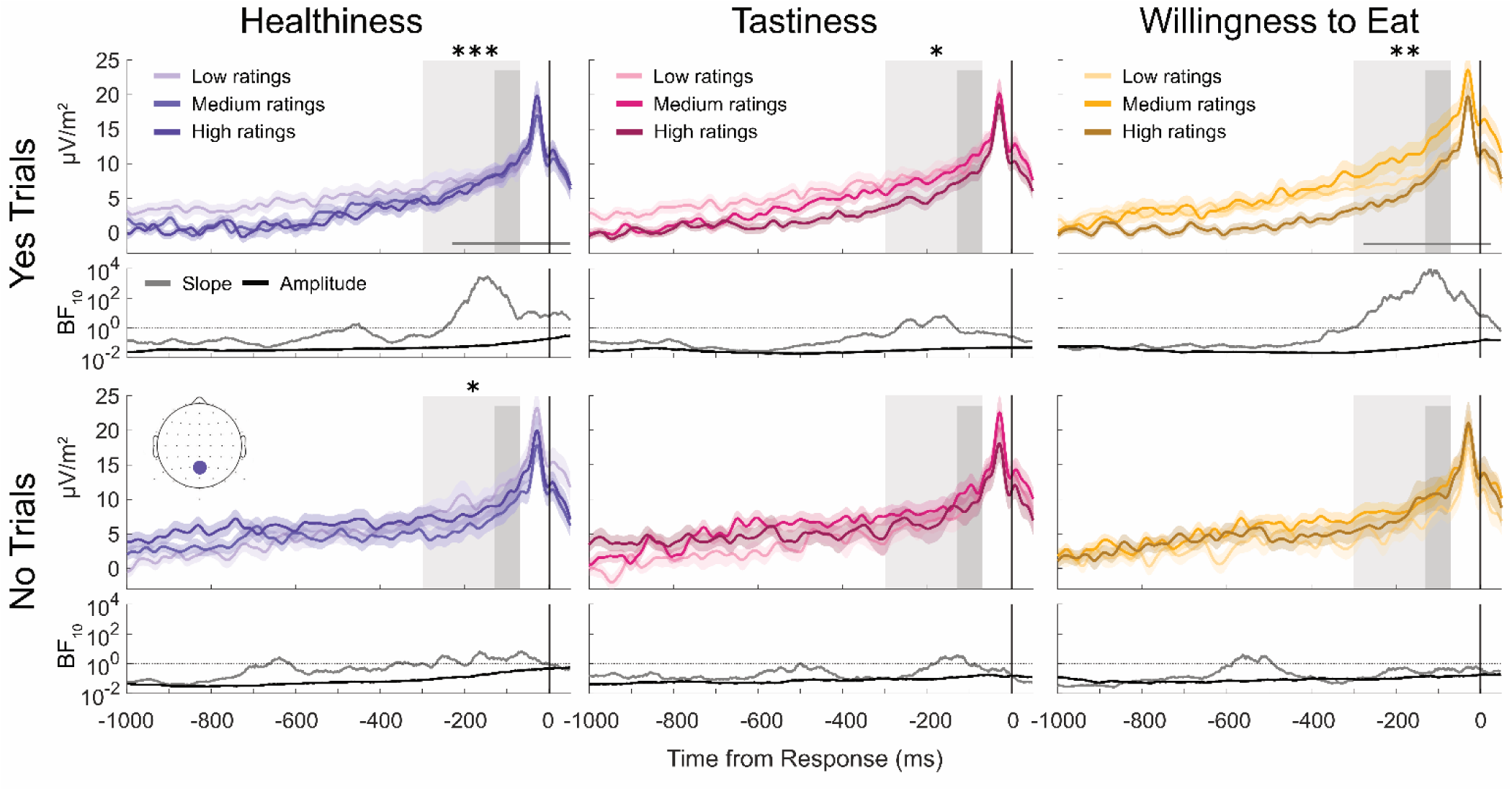
Associations between continuous ratings and CPP slope and amplitude. Group-averaged response-locked ERP waveforms for Yes and No trials in healthiness, tastiness, and willingness to eat decisions, split by attribute rating tertiles. The shaded areas around the plot lines show standard errors of the mean. Pre-response analysis windows are denoted with light grey shading for slope (−300 ms to −70 ms relative to response onset) and dark grey shading for amplitude (−130 ms to −70 ms). Asterisks indicate statistically significant associations between ratings and CPP slope and amplitude within the respective pre-response windows (* indicates *p* < .05, ** indicates *p* < .01, *** indicates *p* < .001). The horizontal grey lines in the upper plots indicate the time windows with statistically significant associations between ratings and CPP slope. In the lower plots, grey and black lines indicate the BF over time for the associations between ratings and CPP slope and amplitude respectively.

We also performed mass-univariate analyses to test for rank correlations between ratings and CPP slopes across the full response-locked epoch. For the Yes trials in healthiness and willingness to eat decisions, ratings were significantly correlated with CPP slopes in a window that overlapped with the pre-response analysis window, as denoted by horizontal grey lines in the upper plots (healthiness: from -231 ms to +49 ms, willingness to eat: -278 ms to +25 ms; Figure 4). Within these windows, ratings were positively correlated with CPP slopes, indicating that stronger representations were associated with steeper slopes.

The less consistent slope effects for No trials compared to Yes trials may partly arise from reduced measurement precision due to relatively fewer observed No responses. Across participants, Yes responses were more frequent than No responses for all three decision types, with the discrepancy being more pronounced for tastiness (Yes trials: mean number of trials = 75, *SD* = 13, range = 37-102; No trials: mean = 38, *SD* = 13, range = 2-73) and willingness to eat decisions (Yes trials: mean = 73, *SD* = 13, range = 40-100; No trials: mean = 41, *SD* = 14, range = 5-73). By comparison, healthiness decisions were more balanced between Yes and No responses (Yes trials: mean = 62, *SD* = 62, range = 44-93; No trials: mean = 52, *SD* = 10, range = 19-72). The rating-dependent slope effects for No trials in tastiness and willingness to eat decisions should therefore be interpreted with caution, as reduced trial counts in these conditions would attenuate our ability to detect effects of comparable magnitude to those observed for Yes trials. Consistent with this interpretation, we found a statistically significant negative correlation between continuous ratings and CPP slope for No trials in healthiness decisions within the pre-response analysis window.

We did not find evidence that pre-response CPP amplitudes were associated with subjective attribute values, consistent with an evidence accumulation trajectory to a fixed decision threshold. We did not observe statistically significant associations (Kendall’s tau-b) between CPP amplitudes in the pre-response window and ratings for any of the three decision types. Mass-univariate results did not reveal any statistically significant clusters of correlations between CPP amplitudes and ratings across the response-locked epoch for all decision types.

## Discussion

The CPP is posited as a neural correlate of evidence accumulation in perceptual decisions, exhibiting two key characteristics: steeper build-up rates that are associated with faster RTs and higher stimulus quality (reflecting higher drift rates) and pre-response amplitudes that remain relatively constant across trials (reflecting a fixed decision boundary). We tested whether this neural signature is also observed during dietary decisions. Our task design also allowed us to disentangle neural correlates of decision formation from those underlying attribute appraisal processes.

In our large (N = 110) sample, we observed steeper pre-response CPP slopes in trials with faster RTs across all three decision types (healthiness, tastiness, and willingness to eat). Although pre-response amplitudes were smaller (i.e., less positive-going) for trials with fast RTs, this appears to be a consequence of our deconvolution method. In addition, we found that patterns of categorical choice RTs differed across lower and higher continuous attribute rating trials in a manner consistent with drift rate changes. CPP slopes were also steeper when ratings more strongly accorded with participants’ categorical choices for Yes responses, congruent with changes in drift rate. We did not observe evidence that pre-response amplitudes covaried with continuous ratings for any decision type, consistent with a fixed decision boundary. Taken together, our findings provide converging evidence that the CPP traces the path of evidence accumulation during multiple types of dietary decisions.

These findings further support the extension of evidence accumulation frameworks to investigate the neural processes underlying dietary choice. While substantial research has examined attribute appraisal and subjective value computation (Chae et al., 2026; Hare et al., 2009; Meule et al., 2013; Moerel et al., 2024; Schubert et al., 2021; Toepel et al., 2009), the neural mechanisms that transform these value representations into choices and motor responses have remained largely unexamined. Sequential sampling models provide a computational account of this transformation through noisy evidence accumulation toward a decision boundary in perceptual decisions (Ratcliff & McKoon, 2008; Ratcliff & Smith, 2004), and our findings support the idea that this framework can be extended to a wide range of decision types (Kable & Glimcher, 2009; Shadlen & Shohamy, 2016). Our findings establish that the CPP, a neural marker of evidence accumulation in multiple decision domains (Fong et al., 2026; Pisauro et al., 2017; van Ede & Nobre, 2024), exhibits the same accumulation-to-bound dynamics during dietary decisions.

Previous work has identified neural signatures of evidence accumulation when participants choose between two food items in a value-based comparative judgment task (Pisauro et al., 2017). We make two key extensions to this work. First, we demonstrate that the same accumulation dynamics may underlie decisions about single food items (i.e., choosing whether to eat a specific food). Second, we show that the CPP also traces decision processes for attribute judgements, during which participants evaluate individual foods on healthiness or tastiness rather than making consumption decisions. These extensions are theoretically important, as single-item and attribute-based decisions differ fundamentally from binary choice tasks. When choosing between two foods (e.g., an avocado versus a tomato), participants compare relative subjective values, potentially considering multiple attributes weighted according to importance depending on the specific options presented. In contrast, single-item willingness-to-eat decisions require comparing a food’s subjective value against an internal criterion, which is a structurally different evaluation (e.g., one might choose an avocado over a tomato, yet decline to eat the avocado when it is presented in isolation). Similarly, attribute judgements require evaluating each food on a consistent, specified dimension (healthiness or tastiness) rather than a combination of subjectively important features.

Despite these fundamental differences in decision structure, we found similar accumulation dynamics as reflected in the CPP waveforms across all three decision types. This suggests that the CPP indexes a domain-general evidence accumulation process towards a decision outcome threshold, invariant to whether the evidence constitutes relative comparisons (choice between multiple options), absolute evaluations (single-item choice), or dimension-specific judgements, or whether the source of the evidence is exogenous (sensory) or endogenous (preferences, knowledge, or memory). Our findings support the CPP as a neural correlate of a domain-general decision signal and suggest that diverse forms of decision-relevant information are translated into action via similar processes.

We also demonstrate that isolating neural correlates of evidence accumulation requires careful application of signal processing techniques. We applied CSD transformations to minimise the overlap from concurrent motor preparation signals (Kelly & O’Connell, 2013) and used a deconvolution algorithm (Ouyang et al., 2015) to account for temporally overlapping stimulus-locked activity. Consequently, our findings address recent challenges to whether the CPP traces evidence accumulation in non-perceptual decisions. Recently, researchers have claimed that deconvolution eliminates the characteristic RT-slope association in value-based tasks (Frömer et al., 2024). However, it was shown that the deconvolution method used in that study was not appropriate for recovering an accumulation-to-bound signal as specified in sequential sampling models, which is neither strictly stimulus-locked nor response-locked but ramps between stimulus onset until just prior to the response (for further discussion see O’Connell et al., 2025). We therefore used an algorithm that allowed us to identify stimulus-locked activity and subtract this to better isolate such ramping activity (Ouyang et al., 2015).

While this method was better suited for recovering a signal that traces evidence accumulation, our simulation analyses revealed that this approach can introduce a different artefact (see Supplementary Figure 3). As the onset of the ramping CPP component is aligned to stimulus onset, some amplitude variation attributable to the CPP can be misattributed to stimulus-locked activity and subtracted during deconvolution. This attenuation is greater for faster trials, as stimulus- and response-locked components overlap more closely in time, leading to a spurious amplitude-RT effect. This artifact may have obscured the fixed pre-response amplitude predicted by evidence accumulation models. Note that this artifact produces *lower* pre-response amplitudes for faster trials, which is the opposite pattern to the pre-deconvolution findings in the study by Frömer et al. (2024). We consider this a necessary trade-off, as failing to adequately control for stimulus-locked activity risks misattributing overlap effects to accumulation processes, a more fundamental confound than the small amplitude artifact introduced by our deconvolution method. Importantly, we found that the CPP slope was modulated by both RT and continuous attribute ratings after deconvolution, which aligns with predictions of evidence accumulation models and provides support for accumulation processes in dietary decisions despite the amplitude artefact. Future work may develop deconvolution techniques that can better isolate accumulation signals while minimising artefactual amplitude modulation.

Our study used continuous ratings as a measure of subjective attribute value and found that ratings covaried with CPP slopes in Yes choice trials in a manner consistent with predictions of evidence accumulation models. This extends previous work demonstrating similar associations in other decision domains. In perceptual decision tasks, stimulus quality has been shown to modulate CPP slope (Feuerriegel et al., 2021; Kelly & O’Connell, 2013; Twomey et al., 2015), while in binary food choices, steeper CPP slopes have been associated with larger estimated value differences between options (Pisauro et al., 2017). Larger perceptual discriminability or value differences indicate higher quality of evidence favouring one option over another in these tasks. We demonstrate that continuous ratings provide a comparable index of subjective attribute value for single-item willingness-to-eat decisions and attribute judgements. Higher ratings for Yes responses and lower ratings for No responses indicate evaluations that are further from one’s subjective categorical boundary, corresponding to stronger representations analogous to higher discriminability in perceptual tasks, whereas middling ratings near the boundary are more ambiguous and reflect weaker evidence.

Consistent with drift rate modulations in evidence accumulation models, CPP build-up rates scaled systematically with subjective attribute value for Yes trials, with steeper slopes for trials with higher ratings. On the other hand, slope modulation by subjective attribute value for No trials were less consistent, with effects only observed in the specified pre-response analysis window for healthiness decisions. This may partly reflect differences in statistical power arising from an imbalance in trial counts between Yes and No responses for tastiness and willingness to eat decisions, potentially due to participants’ general tendency to rate foods positively on these dimensions. The lack of clear evidence for slope effects for the No trials in tastiness and willingness to eat trials should therefore be interpreted cautiously, as reduced trial counts would attenuate our ability to detect effects. Future work may ensure more balanced response distributions to provide sufficient measurement precision (e.g., through careful curation of stimuli). Nevertheless, our finding that continuous CPP slope showed weaker associations with ratings than with RT aligns with predictions from evidence accumulation models.

While drift rate, which we assumed will be higher for subjective attribute values more strongly in favour of one’s choice (Polanía et al., 2014), determines the average rate of accumulation, the evidence accumulation trajectory on any given trial is substantially influenced by within-trial decision noise in the DDM. Single-trial RTs reflect this trial-by-trial variability in both drift rate and noise, whereas continuous ratings index only the systematic drift component. Similar to our results, Pisauro et al. (2017) found that CPP slope modulation by RT was greater than modulation by stimulus value differences. More generally, the relative magnitudes of RT and subjective attribute value associations will depend on the spread of RTs within each condition or subset of trials, compared to the average RT differences between them.

Our findings provide a solid foundation for future joint modelling approaches to formally quantify relationships between CPP dynamics and drift rate by simultaneously fitting behavioural and neural data (Bridwell et al., 2018; Palestro et al., 2018; Sun et al., 2025; Turner et al., 2019). Although a joint modelling venture is beyond the scope of the current study, our finding that CPP slope is modulated by RT and subjective attribute value in theoretically predicted ways forms a strong basis for formally establishing a model linking CPP build-up rates to drift rate parameters. Establishing this neural correlate will enable researchers to develop more comprehensive mechanistic accounts of how subjective value signals or attribute information are converted into action outcomes during dietary decisions.

Our analysis framework can also be applied to investigate how individual differences and contextual factors interact with the underlying computational processes driving food choice. For example, researchers may examine whether individual differences in food preferences, eating motivations, dieting status, or physiological state (e.g., hunger), or the effect of contextual manipulations (e.g., social settings, dietary interventions, or priming) manifest as changes in specific parameters of the DDM. If an intervention alters RT distributions or between-group differences in RTs are observed, joint models incorporating neural measurements can reveal which model parameters correspond to these changes (De Hollander et al., 2016; Turner et al., 2019). For example, changes in drift rate may reflect differences in stimulus quality, subjective attribute value, or attention, while differences in threshold and starting point may indicate differences in caution or response bias (Voss et al., 2004). Computational modelling of choice and RT data can clarify which parameters are affected, and incorporating neural measures provides the mechanistic link, revealing how these computational processes may be implemented by the brain. Explicitly considering evidence accumulation dynamics also affords new perspectives on patterns of speeded dietary choices. The influence of within-trial noise during evidence accumulation is expected to produce a wide range of RTs even for the same food item, meaning that the associations between single-trial RTs and subjective food item evaluations (drift rates) may be weak. Inconsistency across choices made at different times may also reflect the influence of within-trial decision noise rather than true changes in underlying choice preferences.

In conclusion, we demonstrate that the CPP exhibits signatures of evidence accumulation during multiple types of dietary decisions, with CPP build-up rate scaling with RT and subjective attribute value. Our findings indicate that diverse sources of decision-relevant information, from perceptual features to subjective attribute evaluations, are translated into action via shared accumulation mechanisms during dietary decision-making, where this accumulation trajectory is indexed by the morphology of the CPP.

## Data and Code Availability Statement

Data processing and analysis code will be available via the Open Science Framework (https://osf.io/j247x/) at the time of publication. The raw and processed continuous EEG data are available via OpenNeuro (https://doi.org/10.18112/openneuro.ds007012.v1.1.1). For more detail on this experiment and EEG data, see the dataset descriptor paper (Chae et al., 2025).

