## Supplementary Material for "The Centroparietal Positivity Traces the Path of Evidence Accumulation During Dietary Decisions"

Redmond Barry Building,

University of Melbourne, VIC 3010, Australia

CPP Pre-Response Amplitudes by RT Quantile

To visualise the associations between RTs and CPP pre-response amplitudes in a more detailed manner than what is visible in the RT tertile ERP plots, we plotted amplitudes as a function of RT quantile bin. For each participant we divided the data into 20 RT quantiles, each capturing 5% of trials. For example, the first quantile (leftmost data point in each plot) captured the 5% of trials with the fastest RTs. Mean RTs were calculated for each RT quantile and were then averaged at the group level for plotting on the X axes.

As visible in Supplementary Figure 1, pre-response amplitudes were smallest for the fastest RTs and gradually increased in amplitude with increasing RT until around 1000 ms from choice prompt onset. Amplitudes appeared to be relatively stable across quantiles for RTs of 1000 ms onward.

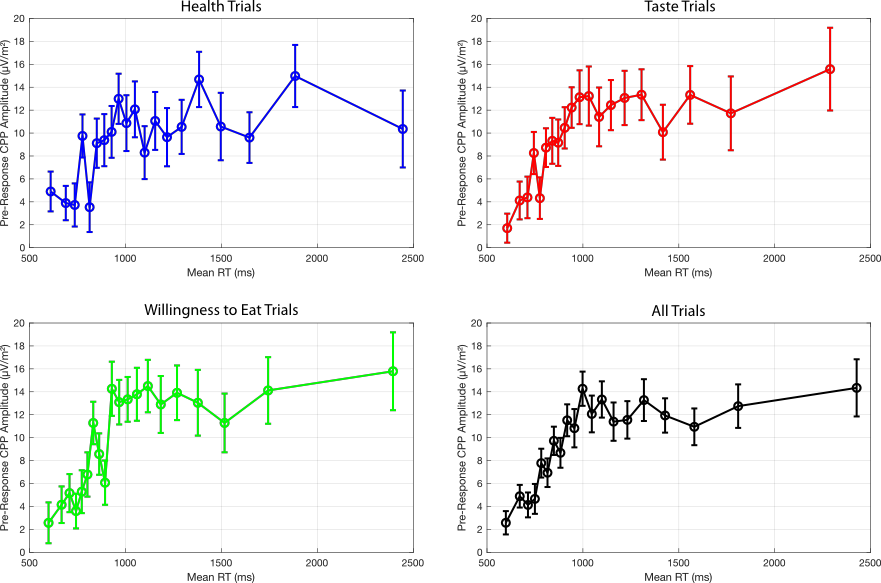

**Supplementary Figure 1. Group averaged CPP pre-response amplitudes by RT quantile bin.** Data points from left to right in each plot denote CSD-transformed CPP amplitudes at electrode Pz for the fastest to slowest RT quantiles, in steps of 5%. The X axis denotes the mean RTs within each quantile bin, averaged across the sample. Data are plotted for each decision type separately and the average of quantile values across decision types. Error bars denote standard errors. Amplitudes appear to gradually increase up to RT values of approximately one second, after which amplitudes are more stable across the range of slower RTs.

Yes/No Continuous Attribute Rating Discrepancies by RT Quantile

We assessed the degree of consistency across categorisation choices and continuous ratings, for categorisation decisions made with different RTs. To do this, we first divided the RT data into quantiles (in bins of 5%, from fastest to slowest). We then calculated the mean continuous rating values for Yes and No trials separately for the food images presented in each set of trials. Difference measures were calculated as the difference in mean ratings between Yes and No choice trials. Large positive values mean that foods that elicited Yes choices for trials within a given RT bin had much higher continuous ratings than those for which No choices were given, indicating strong consistency across categorical and continuous attribute judgment tasks. Low values would conversely indicate less concordance and would suggest some proportion of random responding in the categorical choice task. We assessed whether this may be the case for the fastest RTs (leftmost data points in each plot), analogous to fast guessing responses in perceptual discrete choice tasks. For a small number of participants who did not have any Yes or No trials in a given RT quantile, no difference measure could be calculated and so were excluded from averaging for that quantile. The number of excluded participants per RT quantile is reported in Supplementary Table 1.

Mean rating differences were generally high across both fast and slow RT ranges and were larger for faster as compared to slower RTs, consistent with the observed patterns of correlations between continuous ratings and choice RTs in Figure 1C. There was only a slight drop in the rating difference measures for the fastest RT quantiles, suggesting a low proportion of fast guessing responses in these very fast RT trials.

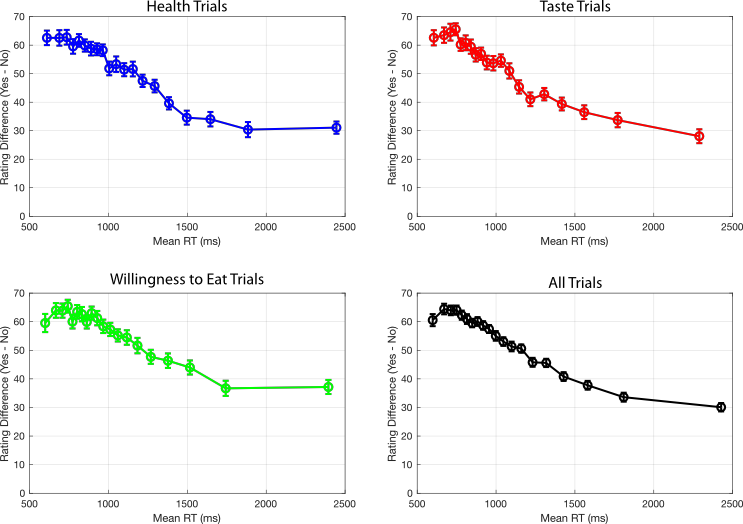

Supplementary Figure 2. Group averaged [Yes – No] continuous attribute rating differences by RT quantile bin. Data points from left to right in each plot denote [Yes-No] mean continuous attribute rating differences for the fastest to slowest RT quantiles, in steps of 5%. The X axis denotes the mean RTs within each quantile bin, averaged across the sample. Data are plotted for each decision type separately and the average of quantile values across decision types. Error bars denote standard errors. [Yes – No] differences appear to be slightly smaller for the fastest RT quantiles in some decisions and become gradually lower for slower RT quantiles.

Supplementary Table 1. Number of excluded participants by RT quantile bin. Participants who did not have any Yes or No trials in a given RT quantile were excluded from averaging for that quantile, as the [Yes – No] difference measure could not be calculated.

| RT percentile bin | Number of excluded participants (/110) | | | |
| --- | --- | --- | --- | --- |
|  | Healthiness | Tastiness | Willingness to eat | All |
| 0-5 | 24 | 34 | 26 | 4 |
| 5-10 | 26 | 38 | 29 | 3 |
| 10-15 | 16 | 35 | 25 | 3 |
| 15-20 | 11 | 27 | 22 | 2 |
| 20-25 | 11 | 34 | 22 | 0 |
| 25-30 | 8 | 27 | 18 | 3 |
| 30-35 | 7 | 20 | 13 | 1 |
| 35-40 | 7 | 25 | 17 | 0 |
| 40-45 | 10 | 17 | 11 | 0 |
| 45-50 | 9 | 15 | 16 | 0 |
| 50-55 | 6 | 14 | 10 | 0 |
| 55-60 | 6 | 14 | 10 | 1 |
| 60-65 | 9 | 9 | 8 | 0 |
| 65-70 | 4 | 12 | 9 | 0 |
| 70-75 | 6 | 13 | 15 | 0 |
| 75-80 | 7 | 16 | 12 | 0 |
| 80-85 | 6 | 8 | 8 | 0 |
| 85-90 | 8 | 9 | 13 | 0 |
| 90-95 | 3 | 11 | 7 | 0 |
| 95-100 | 3 | 8 | 10 | 0 |

**RIDE Deconvolution Simulations Using Artificial Data**

To evaluate whether the observed associations between centroparietal positivity (CPP) pre-response amplitude measures and response times (RTs) may have been an artefact of our deconvolution procedure, we conducted a simulation analysis in which the ground-truth signal was fully known. Similar to simulations of the CPP component by O’Connell et al. (2025), we generated simulated data containing a signal that ramps from a fixed latency after stimulus onset and peaks at a fixed amplitude at the time of response. We assessed whether the Residue Iteration Deconvolution (RIDE) algorithm (Ouyang et al., 2015) systematically underestimates the peak amplitude of the ramping signal for fast RT trials under simulated conditions that are similar to our own dataset.

We generated 2,000 trials of simulated EEG data for a single channel. Each trial consisted of a 3,500 ms epoch (1,000 Hz sample rate) containing two additive signal components. The stimulus-locked component (Supplementary Figure 3B) consisted of a Gaussian-shaped waveform peaking at 500 ms from stimulus onset (peak amplitude = 5 amplitude units, SD = 150 ms). This simulated signal is comparable to the estimated S component waveform at electrode Pz derived from RIDE in the main analyses (Supplementary Figure 4A). A ramping component (Supplementary Figure 3B) consisted of a linearly increasing signal beginning at 500 ms from stimulus onset that rose until the time of the response (peak amplitude = 10 amplitude units). Following the response, the signal linearly decreased back to zero over a fixed 200 ms ramp-down period, mimicking the typical morphology of the CPP component. Please note that this simulated ramping signal cannot be clearly categorised as either stimulus- or response-locked. Supplementary Figure 3E displays the trial-averaged combined signal with both components and Supplementary Figure 3A shows examples of the combined signal from individual simulated trials.

Trial-by-trial RTs were drawn from a gamma distribution (shape parameter = 5, scale = 200 ms, offset = 620 ms) to approximate the right-skewed RT distribution in our data (Supplementary Figure 3F). As the ramping component had a consistent peak amplitude at the time of the response irrespective of RT, the slope of the ramping signal covaried with RT. Steeper slopes occurred in simulated trials with faster RTs, as visible in plots of the ramping component that are time-locked to the response (Supplementary Figure 5A).

The RIDE algorithm was applied to the simulated data to isolate three components: a stimulus-locked (S) component estimated from 1 to 1000 ms relative to stimulus onset, a response-locked (R) component estimated from −600 to 100 ms relative to response onset, and a time-varying (C) component estimated from 500 to 2000 ms relative to stimulus onset. Deconvolution algorithm settings were comparable what was used for our EEG analyses.

We observed that simulated EEG signals originating from ramping activity were partly misattributed to the S component when RIDE was applied. As visible in Supplementary Figure 3C, the estimated S component captured most of the stimulus-locked signal but additionally included a portion of the ramping signals that occurred outside of the stimulus-locked signal time window. Next, we subtracted the estimated S component from each trial of simulated data to recover the ramping signal (Supplementary Figure 3D). Following S component subtraction, data were realigned to the time of the response. The response-locked recovered ramping signal was then compared to the ground-truth ramping signal (Supplementary Figure 5).

We observed an attenuation of ramping signal amplitude that was most pronounced around the time of the response for trials with fast RTs. Amplitude for the fast RT trials was attenuated to a greater degree compared to the medium and slow RT trials in the recovered ramping signal, despite the amplitude at the time of the response being fixed across faster and slower RT trials in the originally simulated ramping signals (Supplementary Figure 5). This attenuation arises because the signals associated with the early portion of the ramp, which begins at a fixed latency post-stimulus, are consistent in time relative to stimulus onset. Consequently, some of this signal is erroneously attributed to the estimated S component when applying the RIDE algorithm. As fast RT trials have shorter intervals between stimulus onset and the response, the amplitude reduction after S component subtraction near the time of the response is consequently larger compared trials with to medium and slow RT trials. For medium and slow RT trials, stimulus onset is temporally more distant from the response, and so the attenuation is smaller or primarily occurs in early time windows relative to the RT. This results in a lower amplitude around the time of the response for fast as compared to slower simulated RT trials.

These simulations demonstrate a specific mechanism by which RIDE deconvolution can produce spurious RT-amplitude associations in ramping signals that are consistent with amplitude variation that we observed in our data. As visible in Supplementary Figure 4, the time-course of the decrement in pre-response CPP amplitudes by RT in our EEG data closely mirrors the profile of the group-averaged S component estimated using RIDE in our dataset. Misattribution of ramping activity to the S component during the 1000 ms post stimulus time window would be expected to reduce pre-response CPP amplitudes in the manner observed in our data.

Notably, the slope differences across RT tertiles were still visible in the RIDE S component-subtracted data. This indicates that RIDE is still a useful method for isolating stimulus-locked from latency-varying EEG signals to characterise evidence accumulation dynamics.

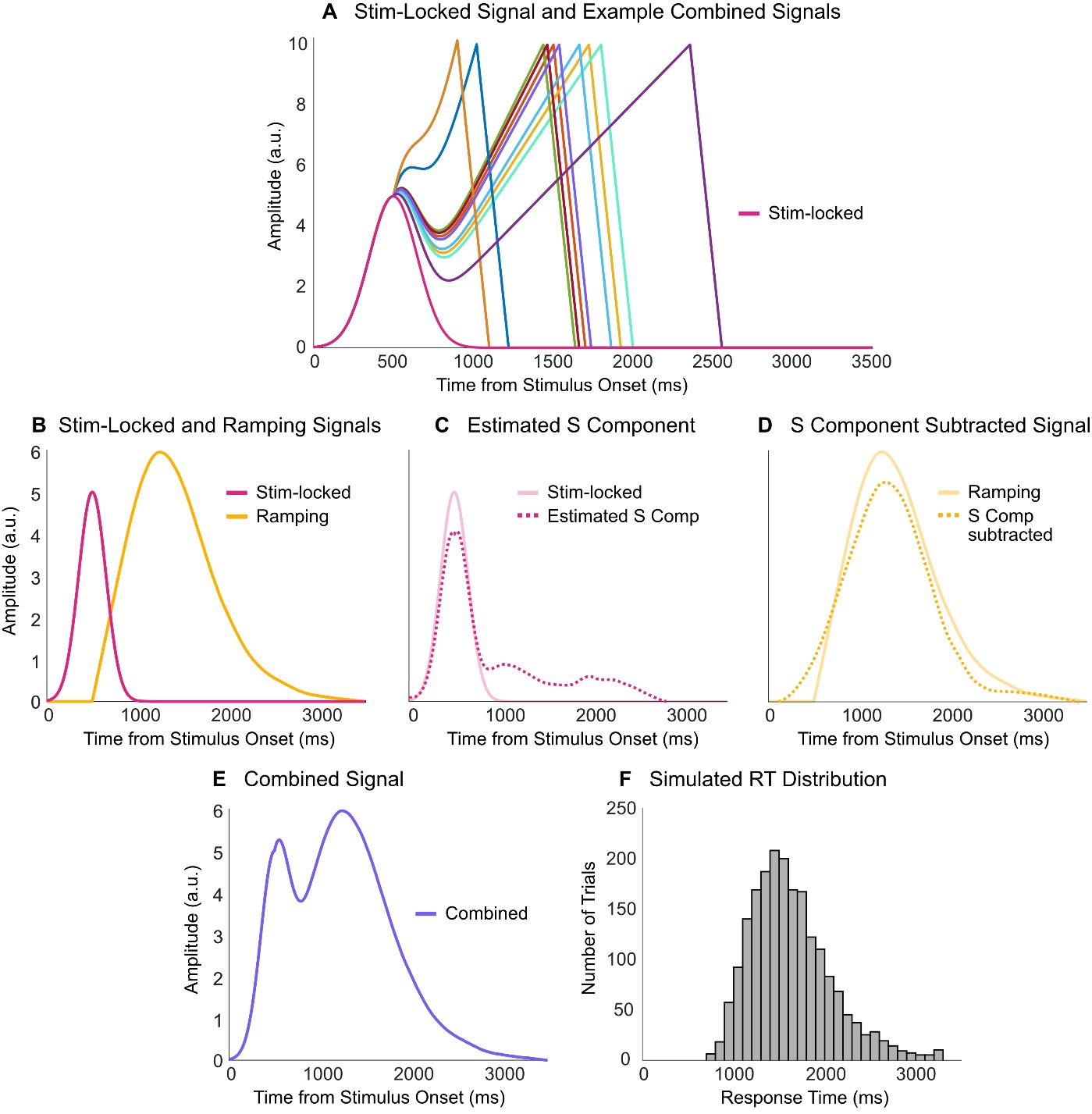
**Supplementary Figure 3. Simulation of stimulus-locked and ramping signals and the simulated RT distribution.** A) Trial-averaged stimulus-locked signal (solid pink) and 10 examples of the simulated signals in single trials with varying RTs (combined signals after adding together the stimulus-locked and ramping signals). B) Trial-averaged plots of the stimulus-locked signal (solid pink; a Gaussian-shaped waveform peaking at 500 ms from stimulus onset) and ramping signal (solid yellow; a linearly increasing signal beginning at 500 ms from stimulus onset and rising until the time of the response to a fixed amplitude). C) Estimated S component after running RIDE deconvolution (dotted pink). Most of the ground-truth stimulus-locked signal (solid light pink) is captured by this component, but some of the ramping signal is also erroneously captured. D) S component-subtracted signal (dotted yellow) mostly captures the ramping signal (solid light yellow). E) Trial-averaged combined signal after adding together the stimulus-locked and ramping signals. F) Simulated response time distribution with a right skew.

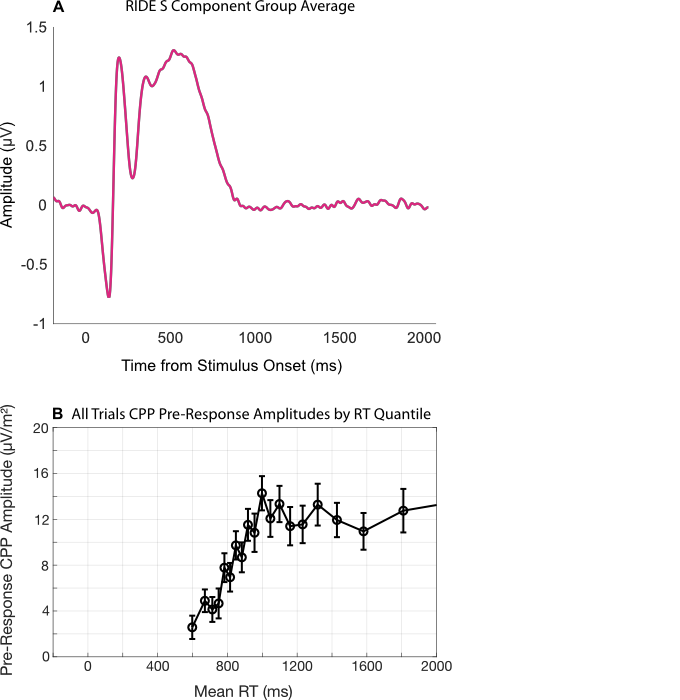

**Supplementary Figure 4. RIDE-derived group-averaged S (stimulus-locked) component waveform at electrode Pz.** A) The estimated S component captures ERP waveform contributions that have consistent timing relative to stimulus onset. B) Group average pre-response CPP amplitudes by RT quantile bin averaged across trial types. Data points from left to right in each plot denote CSD-transformed CPP amplitudes at electrode Pz for the fastest to slowest RT quantiles, in steps of 5%. The X axis denotes the mean RTs within each quantile bin, averaged across the sample. Error bars denote standard errors. Note that the X axis range has been standardised across both plots. The observed pre-response amplitude profile appears to mirror the S component estimate, with largest amplitude decrements when the S component is most prominent. Please note that Y axis units are in µV for the RIDE S component, which are much smaller than the µV/m² units used for CSD-transformed CPP amplitudes.

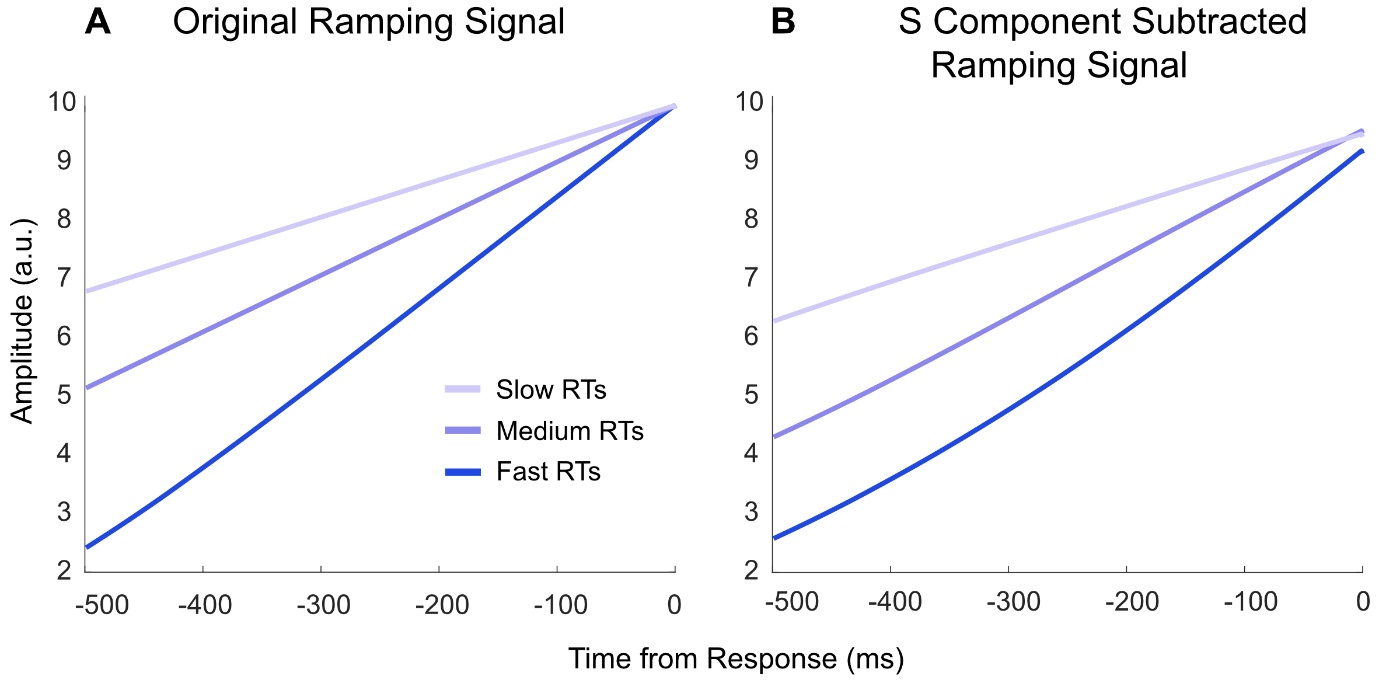
**Supplementary Figure 5. Effect of the RIDE deconvolution procedure on the ramping signal.** A) Original ramping signals averaged over trials within fast, medium and slower RT tertile bins. B) S component subtracted data time-locked to the response. The RIDE deconvolution visibly attenuates the amplitude at the time of the response for the fast relative to medium and slow RT trials, consistent with the patterns of EEG data we observed. The modulation of slope by RT is largely preserved after deconvolution.

**Linear Mixed Effects Model Equations and Coefficients**

We fitted linear mixed-effects models (LMMs) using the lme4 package (Bates et al., 2015) to test whether RT predicted CPP slope and amplitude separately for each decision type (healthiness, tastiness, willingness to eat). RT was log-transformed and then z-scored within each participant. We specified a maximal random-effects structure that included participant intercepts and random slopes for the predictor of interest. If the maximal model produced a singular fit or failed to converge, we removed the random slope and retained the random intercept.

Models were compared with and without the fixed effect of RT (i.e., m4 vs m3, or m2 vs m1) using maximum likelihood and Chi-square likelihood-ratio tests with matched random-effects structures across models. Likelihood ratio test statistics are reported in Supplementary Table 2 and regression coefficients are reported in Supplementary Table 3. Models are denoted in lme4 notation as follows:

m1: *CPP_measure_* ∼ 1 + (1 | *pID*)

m2: *CPP_measure_* ∼ 1 + *RT* + (1 | *pID*)

m3: *CPP_measure_* ∼ 1 + (1 + *RT* | *pID*)

m4: *CPP_measure_* ∼ 1 + *RT* + (1 + *RT* | *pID*)

**Supplementary Table 2. Chi-square likelihood ratio tests for group-level model comparisons.**

| Decision type | CPP measure | Chi-square | df | p | Comparison test | Best model |
| --- | --- | --- | --- | --- | --- | --- |
| Healthiness | Slope | 48.46 | 1 | <.001 | m1 vs m2 | m4 |
|  |  | – | – | – | m3 vs m4 |  |
|  | Amplitude | 19.21 | 1 | <.001 | m1 vs m2 | m4 |
|  |  | 11.09 | 1 | <.001 | m3 vs m4 |  |
| Tastiness | Slope | 18.40 | 1 | <.001 | m1 vs m2 | m4 |
|  |  | 14.21 | 1 | <.001 | m3 vs m4 |  |
|  | Amplitude | 36.47 | 1 | <.001 | m1 vs m2 | m4 |
|  |  | 12.02 | 1 | <.001 | m3 vs m4 |  |
| Willingness to eat | Slope | 24.72 | 1 | <.001 | m1 vs m2 | m2 |
|  |  | – | – | – | m3 vs m4 |  |
|  | Amplitude | 48.32 | 1 | <.001 | m1 vs m2 | m4 |
|  |  | 15.76 | 1 | <.001 | m3 vs m4 |  |

*Note*. m1: null model, m2: random intercept model, m3: null random slope model, m4: random slope model. Models denoted in red produced a singular fit and failed to converge. Dashes indicate data were not reported as one or more models in the comparison test failed to converge.

**Supplementary Table 3. LLM coefficients for predicting CPP pre-response amplitudes and slopes from RTs.**

| Decision type | CPP measure | Model | Estimate | SE | t | p |
| --- | --- | --- | --- | --- | --- | --- |
| Healthiness | Slope | m2 | -0.02 | <0.01 | -6.97 | <.001 |
|  |  | m4 | -0.02 | <0.01 | -5.68 | <.001 |
|  | Amplitude | m2 | 2.10 | 0.48 | 4.39 | <.001 |
|  |  | m4 | 2.14 | 0.63 | 2.41 | <.001 |
| Tastiness | Slope | m2 | -0.01 | <0.01 | -4.29 | <.001 |
|  |  | m4 | -0.01 | <0.01 | -3.88 | <.001 |
|  | Amplitude | m2 | 2.89 | 0.48 | 6.05 | <.001 |
|  |  | m4 | 2.85 | 0.80 | 3.57 | <.001 |
| Willingness to eat | Slope | m2 | -0.01 | <0.01 | -4.97 | <.001 |
|  |  | m4 | – | – | – | – |
|  | Amplitude | m2 | 3.30 | 0.47 | 6.96 | <.001 |
|  |  | m4 | 3.26 | 0.79 | 4.12 | <.001 |

*Note*. m2: random intercept model, m4: random slope model. Models denoted in red produced a singular fit and failed to converge. Dashes indicate data were not reported as the model failed to converge.
